# Comparative kinetic analysis of ascorbate (Vitamin-C) recycling dehydroascorbate reductases from plant and human

**DOI:** 10.1101/2021.08.28.458007

**Authors:** Bhaba Krishna Das, Amit Kumar, Sreeshma Nellootil Sreekumar, Kannapiran Ponraj, Kaustubh Gadave, Saravanan Kumar, V. Mohan Murali Achary, Pratima Ray, Malireddy K. Reddy, Arulandu Arockiasamy

**Author notes:** Correspondence should be addressed to **Correspondence:** Arulandu Arockiasamy, Membrane Protein Biology Group, International Centre for Genetic Engineering and Biotechnology, Aruna Asaf Ali Marg, New Delhi-110067. India., Ext-172, Alternate. these authors equally contributed to this work (Bhaba Krishna Das, Amit Kumar, Sreeshma Nellootil Sreekumar, Kannapiran Ponraj).

## Abstract

Ascorbate, a primary antioxidant, gets readily oxidized to dehydroascorbate (DHA). Hence, recycling by dehydroascorbate reductase (DHAR) enzymes is vital for protection from cellular oxidative stress in eukaryotes. However, a detailed kinetic analysis of plant DHARs and their human orthologs; chloride intracellular channels (*Hs*CLICs) is lacking. We demonstrate that DHAR from stress adapted pearl millet *Pennisetum glaucum* (*Pg*DHAR) shows the highest turnover rate whereas *Hs*CLIC1, 3, and 4 reduce DHA, albeit at lower rates. We further show that the catalytic cysteine is susceptible to varying levels of oxidation, supported by crystal structures and mass-spectrometry analysis. The differences in kinetic parameters among plant and human DHA reductases corroborate with the levels of reactive oxygen species H_2_O_2_ encountered in their respective intracellular environment. Our findings may have broader implications in crop improvement using pearl millet DHAR, and anti-cancer therapeutics targeting Vitamin-C recycling capability of human CLICs.

## Introduction

Ascorbate (AsA) recycling is an essential process in higher eukaryotes and is primarily carried out by dehydroascorbate (DHA) reductases. In plants, both dehydroascorbate reductase (DHAR) and monodehydroascorbate reductase (MDHAR) enzymes of the glutathione-ascorbate pathway recycle AsA from DHA and MDHA, respectively [1-3]. Similarly, a soluble form of human chloride intracellular channels (*Hs*CLICs) that share high structural homology with plant DHARs is shown to recycle ascorbate from DHA [4]. Both plant DHARs and *Hs*CLICs belong to the glutathione-S-transferase (GST) - Omega family of proteins [5]. DHAR along with MDHAR diligently maintains the AsA pool during conditions of oxidative stress [6, 7] making AsA one of the major low molecular weight antioxidants in plants, in addition to GSH [8]. AsA concentration in plants ranges from 20 to 50 mM [5, 9]. DHA is reduced to AsA by DHAR using GSH as an electron donor [10]. AsA regeneration is vital for maintaining redox homeostasis within the cell [11] and is found to be higher under drought conditions [12]. Further, increased transcript levels of DHAR genes, DHAR activity and AsA recycling were observed under oxidative stress [6]. Arabidopsis mutants devoid of cytosolic DHAR show lower levels of apoplastic AsA and increased ozone sensitivity [13]. Overexpression of *Arabidopsis thaliana* cytosolic DHAR in tobacco results in increased tolerance to drought, ozone and aluminium induced stress. DHAR overexpression in tomato results in better tolerance to methyl viologen and salt stress with a corresponding increase in cellular AsA pool [14]. Overexpressing of DHAR in rice increases AsA pool and crop yield [15]. Therefore, AsA recycling capability imposes a major impact upon the environmental adaptability of plants and also on their yield.

Phylogenetic analysis of pant DHARs and human chloride intracellular channels (CLICs) have long suggested shared functional orthology [16]. *Hs*CLICs exist in both soluble and membrane integrated ion channel forms. *Hs*CLICs are associated with various cellular processes such as tubulogenesis [17], apoptosis [18], cell cycle regulation [19], and redox sensing in humans [20]. The human genome encodes 6 CLICs (*Hs*CLIC1-6), with known splice variants. Though plant DHARs show low sequence identity with *Hs*CLICs (≤ 30 %) they share high structural similarity with a RMSD of ≤ 1.9 Å (Table S1). DHA reductase activity for the soluble form of CLICs was demonstrated [4, 21], owing to the presence of a redox-sensitive CXXC[S/A] motif [16] using indirect assays. Further, oxidoreductase activity of *Hs*CLIC3 [21] is shown to be associated with migration in cancer cells and tumor metastasis, suggesting that Vitamin-C recycling by *Hs*CLICs could be novel drug targets.

Notably, plant DHARs and *Hs*CLICs have a conserved catalytic cysteine, a GSH binding G-site and substrate/product binding H-site. This led us to explore the differences between plant DHAR and *Hs*CLICs in terms of ascorbate recycling capability for gaining deeper insights into their functional implications.

In this study, we have carried out a thorough comparative enzyme kinetic analysis using *Pennisetum glaucum* DHAR (*Pg*DHAR) and human CLIC1, 3 and 4. Further, we have investigated the role of active site residues of *Pg*DHAR and *Hs*CLIC1, 3 and 4 using chemical modifications, site-directed mutagenesis, LC-MS/MS and X-ray crystallography followed by enzyme kinetics. We found that the specific activity of plant DHAR is ∼1000 fold higher than the human CLICs, suggesting a possible role of intracellular ROS levels in shaping up the ascorbate recycling rates of these respective DHA reductases.

## Materials and Methods

### Cloning, Overexpression and Purification of *Hs*CLIC1, 3 and 4

Sequence verified cDNA clones for human CLIC1, CLIC3, and CLIC4 were obtained from ATCC (Supplementary Table 1). The protocols for cloning, overexpression and purification of *Hs*CLIC1, 3 and 4 were same as described below. The genes were sub-cloned into pETM30 under NcoI and BamHI sites, with a TEV cleavable N-terminal His_6_-GST tag. Expression was optimized in BL21 (DE3) cells at 20°C in AIM media for 18-24 hours. Cells harvested after culture were lysed by sonication in lysis buffer 50 mM Tris-HCl pH 8.0, 300 mM NaCl, 10 mM Imidazole, 1 mM Benzamidine, 1 mM Phenylmethanesulfonylfluoride fluoride (PMSF), and 3 mM β-ME. The cell lysate was spun at 15,000 g for 45 minutes at 4 °C, and the supernatant was filtered through a 0.22 μm filter. The clarified supernatant was loaded onto a pre-equilibrated 5ml HisTrap FF column (GE), charged with Co^2+^ or Ni^2+^. Upon binding, the column was washed extensively with wash buffer containing 25 mM Tris-HCl pH 8.0, 300 mM NaCl, 10 mM Imidazole and 3 mM β-ME, eluted using a 10-500 mM Imidazole gradient. The peak fractions were pooled, dialyzed, and subjected to TEV cleavage at 4 °C in 10 mM Tris-HCl pH 8.0, 150 mM NaCl, 0.5-1 mM EDTA and 2.0 mM DTT. After dialysis or desalting to remove DTT, His6-GST tag-cleaved protein was obtained in the flow-through of the second round of HisTrap FF column (GE), charged with Co^2+^ or Ni^2+^. Pooled and concentrated protein was passed through Superdex S75 16/60 column (GE) pre-equilibrated with 10 mM Tris-HCl pH 8.0 and 10 mM NaCl with or without 2 mM DTT, concentrated to 10-25 mg/ml before further use. Protein concentrations were estimated using OD280 nm measurement and theoretical molar extinction coefficient (ProtParam/ExPASy).

### Enzyme assays and kinetics

DHA reductase activity of *Pg*DHAR, *Hs*CLIC 1, 3 and 4 were assayed as described by Stahl [9] with minor modifications using both single cuvette and 96 well microplates. The reaction was initiated by adding 2 mM GSH into the 500 μl (single cuvette) or 200 μl (microplate) reaction mixture containing the protein in 50 mM Phosphate (pH 6.8), 0.2 mM EDTA and 0.1 mM DHA. Absorbance of AsA was monitored at 265 nm. Buffer, denatured protein and BSA were independently used as controls for the assay. Curve fitting and values of V_max_ and K_m_ were calculated by non-linear regression function using GraphPad Prism (v5.0 GraphPad Software, San Diego California, USA.

### Mass-spectrometry

25-50 μg of purified or chemically modified *Pg*DHAR and *Hs*CLIC1 were dialyzed against 25 mM Ammonium bicarbonate. Samples were thereafter denatured and subjected to proteolysis by trypsin (Promega) at 37ºC overnight. The reaction was stopped by adding 0.1 % trifluoroacetic acid, and the samples were desalted using a C18 micro spin column (Sartorius), vacuum dried and subjected to MALDI TOF/TOF based analysis. The desalted tryptic peptides were redissolved in 10ul of 0.1%TFA. Equal volumes of sample and α-cyano -4-hydroxycinnamic acid (HCCA) matrix (10 mg/ml) were spotted onto MALDI ground steel target plate for further analysis. Measurements were performed using an Ultraflex III MALDI TOF/ TOF instrument (Bruker Daltonics, Germany), in the positive ion reflector mode. The instrument parameters for acquisition and analysis were set as described earlier [22]. The final mass spectra were produced by averaging 1500 laser shots. The Peptide Mass Fingerprint (PMF) pattern of the tryptic peptides was further annotated using Flexanalysis software (version 3.0). For MS/MS-based sequencing and identification of protein from the peptide, the parameters as reported earlier [22] were used during acquisition and analysis. Briefly, the precursor peptide ions were fragmented in positive mode using the LIFT.lft method (Bruker Daltonics, Germany). Fragmented peptides were analysed and annotated using Flexanalysis (version 3.0) and were subjected to database search using Biotools (version 3.2). The database search parameters were: enzyme – Trypsin; taxonomy - unrestricted; fixed modifications carbamidomethyl (C); variable modifications –oxidation (M); no restriction on protein mass; allowed up to 1 missed cleavage. The fragment masses (tandem mass spectrum) were searched in NCBInr database with MS tolerance of up to +100 ppm and MS/MS tolerance of up to + 0.7da.

### Chemical modification and circular dichroism spectroscopy

*Pg*DHAR and *Hs*CLIC1 were incubated with 1-2 mM Iodoacetamide or Dimedone (Sigma) at 4°C for 1-2 hours. Unbound chemicals were removed by desalting through a G25 spin column or by buffer exchange using a 3kDa ultracel centrifugal device (Amicon) or by dialysis. To ascertain the structural stability of modified proteins, CD spectra for 3.5-5 nM of chemically modified protein was recorded at 190-280 nm, using Jasco J-810 spectropolarimeter in 50 mM phosphate buffer (pH 6.8).

### Mutagenesis

Mutagenic PCR was used to introduce point mutations in both *Pg*DHAR and *Hs*CLIC1 (Table S2) following manufacturers protocol (Stratagene). 2-5μl of the PCR product was used to transform *E. coli* DH5α cells. Positive clones were confirmed by DNA sequencing (Macrogen) Expression and purification of mutants carried out in a similar way as *Pg*DHAR and *Hs*CLICs.

### Crystallization and structure determination

Crystals were grown using sitting drop vapour diffusion method. Prior to setting up plate, protein was treated with 2 mM H_2_O_2_. Crystals were grown in; *Pg*DHAR-Oxy: 0.1 M Sodium Acetate trihydrate (pH 4.6) and 2.0 M Ammonium sulphate and *Hs*CLIC1-Oxy: 0.02 M Sodium potassium phosphate, 0.1 M Bis-Tris Propane (pH 6.5) and 20 % PEG 3350. ∼20 % glycerol was used as a cryo-protectant. Data were collected at 100 K on two different X-ray sources: (i) Rigaku MicroMax-007 (Cu-Kα 1.5418 Å), attached with Mar345 image plate for *Pg*DHAR-oxy (ii) Rigaku FR-E SuperBright rotating anode generator (Cu-Kα 1.5418 Å) attached with R-axis IV++ image plate for *Hs*CLIC1-oxy. Data sets were processed HKL2000 [23] and autoPROC [24], respectively, for *Pg*DHAR-Oxy and *Hs*CLIC1-Oxy. Phases for *Pg*DHAR-Oxy were obtained using BALBES [25] with native *Pg*DHAR (5EVO) as template. For *Hs*CLIC1, molecular replacement was done using native CLIC1 (1K0M) as template in AutoMR module of Phenix suite [26]. Iterative manual model building and refinement were done using COOT [27] and Refmac5 [28], respectively. CSO (Sulphenic acid), CSD (Sulfinic Acid), OCS (Sulfonic acid) form of cysteine were taken from COOT library. Figures were made using PyMol (Schrödinger, Licensed to ICGEB).

## Results and Discussion

### *Pg*DHAR and *Hs*CLICs share conserved active site

First, the crystal structure of *Pg*DHAR was solved to understand the ligand-binding sites in detail [29]. Both plant DHARs and *Hs*CLICs contain two domains; thioredoxin (TRX)-like N-terminal domain (NTD) of the canonical thioredoxin fold and an α-helical bundle containing C-terminal domain (CTD), typical of Glutathione-S-transferases (GSTs) [30, 31] (Figure 1a). The conserved catalytic cysteine of *Pg*DHAR (C20) [32] and *Hs*CLIC1 (C24) of the monothiol CXXS motif is located in the NTD of these proteins (Figure 1b). Notably, *Pg*DHAR shows the highest sequence identity with other plant DHARs such as *Zea mays* (90%) and *Oryza sativa* (84%) and lowest with green alga *Chlamydomonas reinhardtii* (40%) DHARs (Supplementary Table 2). Pairwise structural superposition of *Pg*DHAR with its plant counterparts show minimal variations (RMSD of ≤2.3Å) (Supplementary Figure 1). High structural and sequence identity among plant DHARs suggests that plants, throughout evolution, have maintained sequence and structural features with minimal deviations, concurrence with the indispensable role of DHARs in the plant kingdom. Similarly, structural homology search of *Pg*DHAR using DALI recovered as best hits (Z ≤28.8) with *Hs*CLICs *viz*., *Hs*CLIC1-5 that share the highest sequence identity of ∼30% and a very high structural similarity of ≤1.7Å RMSD. Thus, sequence and structural conservation among plant DHARs and *Hs*CLICs suggest possible functional conservation between their soluble forms. Further, structure-guided sequence alignment reveals a distinct positively charged pocket shared by these proteins [33] (Figure 1c, d). This pocket is occupied by conserved amino acids distributed around the catalytic CXX[C/S] motif [34]. In addition, the following residues; K8, F22, V60, P61, D73 and S74 which are part of the positively charged pockets of *Pg*DHAR are also conserved in *Hs*CLIC1 as well (K13, F26, L64, P65, D76 and T77) (Figure 1d). Notably, these include residues from the canonical GSH-binding “G-site”, and the hydrophobic “H-site”, suggesting shared orthology in terms of the GSH dependent DHA reductase activity. Thus, we decided to compare their enzyme activities using detailed kinetics.

**Figure 1.**
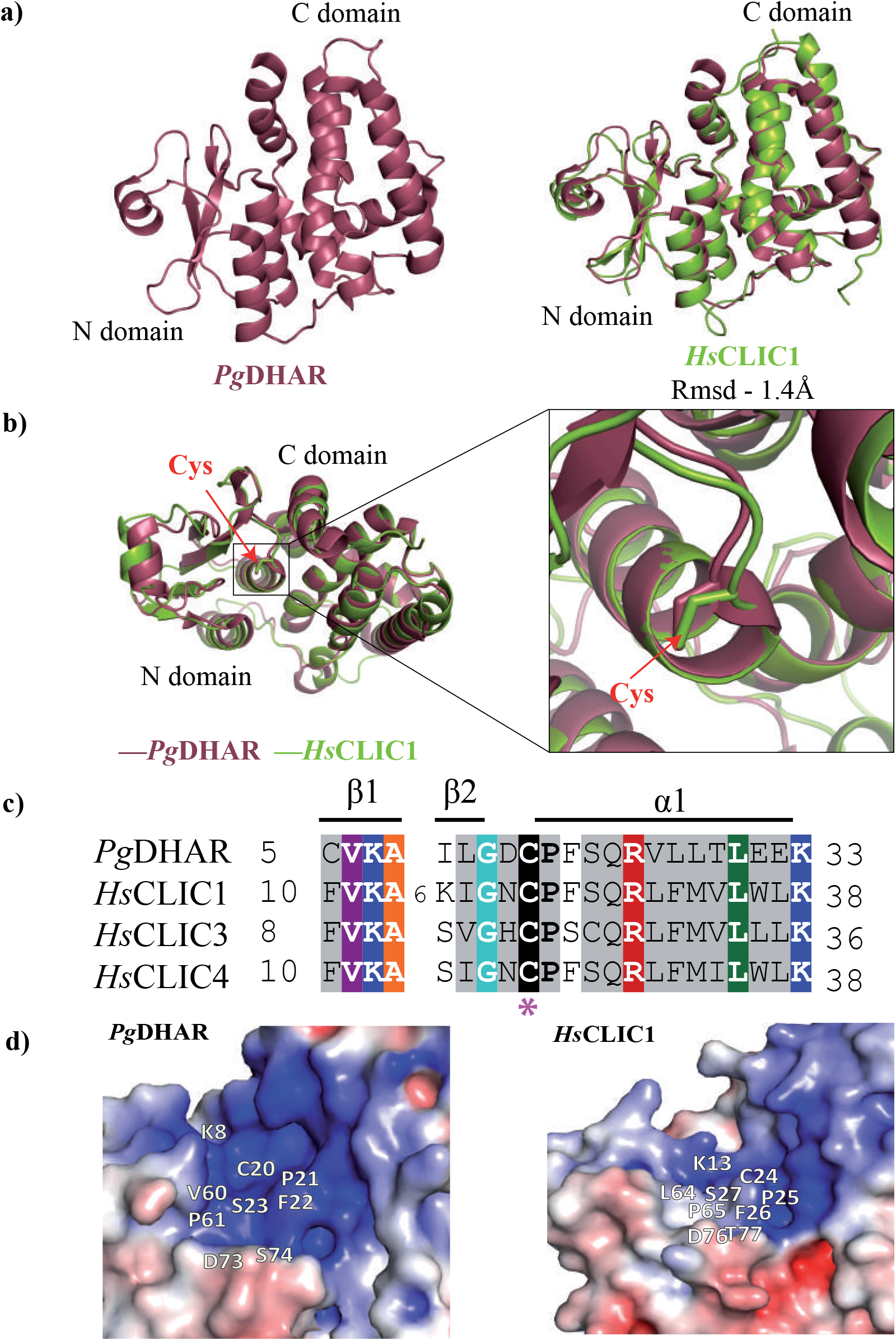
Structural, sequence and active site comparison of *Pg*DHAR with *Hs*CLICs. Structural superposition of (**a**) *Pg*DHAR with *Hs*CLIC1 are shown. Proteins are colour coded as: *Pg*DHAR (5IQY) in raspberry and *Hs*CLIC1 (1K0N) in split pea. **(b)** Ribbon diagram showing the structural superposition of *Pg*DHAR with *Hs*CLIC1. The N-terminal catalytic cysteine in both is shown as a stick model and is zoomed. **(c)** Structure-guided multiple sequence alignment of *Pg*DHAR with *Hs*CLIC 1,3 and 4 showing the conserved CXXS[C/A] motif. Active site cysteine is marked with asterisk (*). **(d)** Putative enzymatic active site of *Pg*DHAR and *Hs*CLIC1 with the conserved residues marked. Protein sequences are labelled with their corresponding protein name, with italic prefix denoting species, and coloured as per the scheme used for structural superposition. The alignment is coloured at 90% consensus using the following scheme: hydrophobic (ACFGHILMTVWY), aliphatic (ILV), aromatic (FHWY), polar (CDEHKQRST), small (ACDGNPSTV), tiny (AGS), big (EFHIKLMQRWY), charged (DEHKR), negative (DE), Ser/Thr (ST), positive (HKR) residues shaded grey; absolutely conserved residues shaded black (**C**); lysine (**K**) residue shaded blue; leucine (**L**) residue shaded green; alanine (**A**) residue shaded orange; arginine (**R**) residue shaded red; glycine (G) residue shaded green. Species abbreviations are as follows: *Pg*: *Pennisetum glaucum, Hs*: *Homo sapiens*.

### *Pg*DHAR shows the highest specific activity and turnover rate

#### 1. *Pg*DHAR *vs* other plant DHARs

Notably, DHAR catalyse the conversion of DHA to ascorbate in a GSH-dependent manner

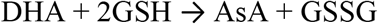

First, we compared the specific activity and turnover rates (k_cat_) of *Pg*DHAR with other known plant DHARs reported in the literature. To do so, recombinant *Pg*DHAR was purified to homogeneity following the published protocol [29]. Using the tag-cleaved protein, detailed enzyme kinetics was carried out using direct assay with DHA and GSH as substrate and cofactor, respectively. *Pg*DHAR follows Michaelis–Menten kinetics, converting DHA to AsA in a GSH-dependent manner. The pH optimum of these enzymes vary considerably; 6.8-7.0 (*Pg*DHAR), 8.2 (*Os*DHAR) [35] and 7.9 (*So*DHAR) [36] (Figure 2a). The reported pH optimum of both *Os*DHAR and *So*DHAR are more towards the alkaline. It is worthwhile to note that DHA auto converts to AsA at higher pH [37] and thus is challenging to distinguish enzymatic conversion versus auto conversion. Kinetic measurements of tag-cleaved *Pg*DHAR yielded K_m_ of 159.2 ±11.4μM and 1477±89.9μM for DHA and GSH, respectively, with a specific activity of 1164 μM min^-1^mg^-1^ and k_cat_ for DHA of 136299.8 min^-1^ (Table 1). *Pg*DHAR shows greater affinity for DHA (159.2 ±11.4μM) than to GSH (1477±89.9μM) when compared to *Os*DHAR (Supplementary Table 3). Of note, when the GSH concentration is kept constant at 2 mM, the affinity (K_m)_ for DHA was roughly 10 times higher than for GSH. Interestingly, the K_m_ and specific activity of tag-cleaved purified *Pg*DHAR reported in this study are different from the one with partially purified *Pg*DHAR containing N-terminal His_6_-tag [32] (Supplementary Table 3).

**Table 1.**
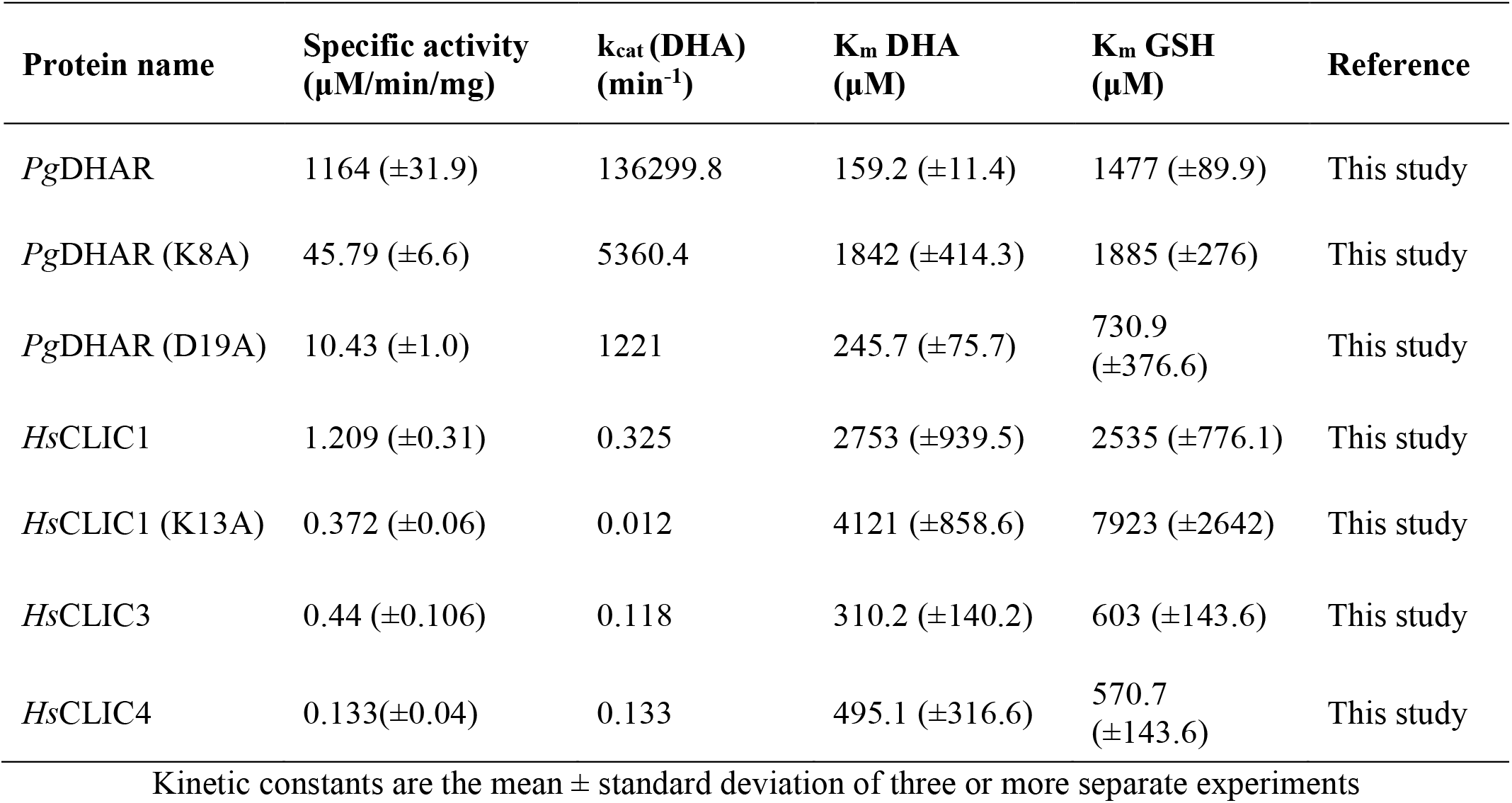
Enzyme kinetics of *Pg*DHAR and human CLICs and their mutants.

**Figure 2.**
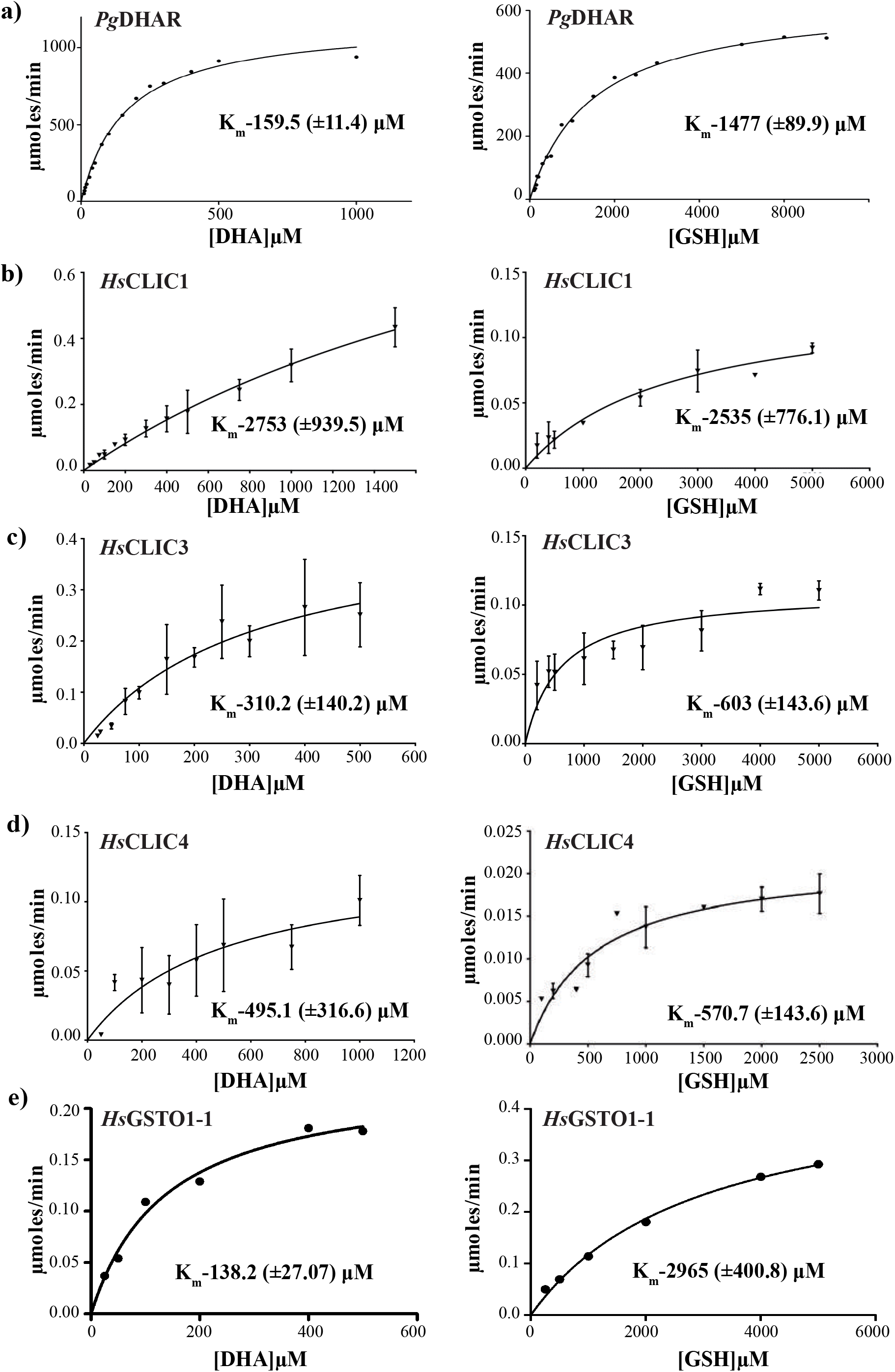
Enzyme kinetics of *Pg*DHAR and *Hs*CLICs. Michaelis-Menten kinetics plot for **(a)** *Pg*DHAR, **(b-e)** *Hs*CLIC1,3,4 and *Hs*GSTO-1 showing DHA reductase activity. The amount of AsA produced per minute under different concentrations of DHA (μM) and GSH (μM) is shown here.

The K_m_ values of all native plant DHARs characterized to date ranges from 0.05 to 0.7 mM and 0.74 to 10 mM for DHA and GSH, respectively (Supplementary Table 3) [38]. Despite the strong sequence, structural and active-site similarities, the specific activity and k_cat_ of *Pg*DHAR were found to be the highest when compared to other plant DHARs. *Pg*DHAR has ∼23 and ∼60-fold increase in specific activity when compared to *Os*DHAR and *Zm*DHAR. The enhanced DHAR activity of *Pg*DHAR is expected to confer efficient recovery of *Pennisetum glaucum* from biotic and abiotic stress such as drought, high temperature and salinity which results in oxidative stress. This observation corroborates well with the geographical distribution of pearl millet.

#### 2. DHA reductase activity of *Hs*CLIC1, 3, 4 *vs Pg*DHAR

A detailed kinetic analysis of the soluble forms of *Hs*CLICs; 1, 3 and 4 were carried out to compare the DHA reductase activity of *Hs*CLICs with *Pg*DHAR. Recombinant *Hs*CLIC1, 3 and 4 were overexpressed in *E. coli* at 20°C and purified to homogeneity using various chromatographic methods and also tag-cleaved. *Hs*CLICs 1, 3 and 4 follow Michaelis–Menten kinetics similar to *Pg*DHAR (Figure 2b-e). K_m_ for DHA and GSH was measured for all three CLICs, for the first time. The respective K_m_ values and specific activities of *Hs*CLICs 1, 3 and 4 are shown in Table 1. Collectively, our data suggest that the affinity of *Hs*CLICs to DHA and GSH is relatively low compared to *Pg*DHAR. The specific activity and k_cat_ of *Hs*CLIC1, 3 and 4 are 1.2, 0.44, 0.133 μM min^-1^mg^-1^ and 0.325, 0.118 and 0.133 min^-1^respectively. Similar k_cat_ and specific activity were observed with other mammalian enzymes reported to have DHA reductase activity [39] (Supplementary Table 4). Affinity for DHA (K_m_ 2753 μM) and GSH (K_m_ 2535 μM) is low for *Hs*CLIC1 with ∼96% reduction in apparent catalytic activity (45.79 μM min^-1^mg^-1^) when compared to *Pg*DHAR (1164 μM min^-1^mg^-1^). Among *Hs*CLICs, affinity for DHA and GSH is maximal for *Hs*CLIC3 (K_m_ DHA 310.2 μM) and *Hs*CLIC4 (K_m_ GSH 570.7 μM), respectively, whereas *Hs*CLIC1 shows highest specific activity (1.209 μM min^-1^mg^-1^). It is interesting to note that the purified forms of these human enzymes show DHA reductase activity *in vitro* with significant differences in specific activity (∼100-1000 fold) and k_cat_ (∼10^6^ fold). This disparity may be attributed to the varying requirement for antioxidant activity in their respective intracellular environments. Notably, the level of intracellular H_2_O_2_ concentrations in plant cells varies from 1-100 μm [40] whereas it ranges between 0.05-0.5 μM in humans [41]. Next, we probed the role of reactive cysteine in CXXC[S] motif of *Pg*DHAR and *Hs*CLIC1.

### Role of catalytic cysteine and its oxidation states

The reactive thiol-containing catalytic cysteine (S^-^ thiolate anion) of DHARs are susceptible to modification by iodoacetamide (IAA) [42], the latter forming a covalent complex with the reactive cysteine. Additionally, these residues also act as redox sensors, which make them amenable to get oxidized, both reversibly and irreversibly, and reduced under varying conditions of cellular redox state. The initial product of cysteine oxidation is sulfenic acid (-SO), which has been reported to be specifically detected by dimedone. The sulfenic form is reversible by the physiological GSH present inside the cell. Hence, we chemically modified *Pg*DHAR and *Hs*CLIC1 using IAA and dimedone and detected multiple oxidation states using mass spectrometry (Supplementary Table 5 and Supplementary Figure 2). MALDI TOF MS/MS was used to successfully identify the parental and alkylated forms of *Pg*DHAR and *Hs*CLIC1. Upon IAA treatment a 57 Da shift at the active site cysteine was detected indicating a mono-alkylation state in concurrence with previous literature [43]. The susceptibility of the active site cysteine in the purified protein to oxidation (Cys-SO) was evaluated by treating with H_2_O_2_ and then dimedone. The catalytic cysteine residue on treatment with dimedone had a 138 Da increment in its molecular weight indicating its conversion to sulfenic form [44].

DHA reductase activity of the chemically modified proteins, where the catalytic cysteine was either alkylated by IAA or oxidised by dimedone, was measured to study the involvement of catalytic cysteines in the reaction mechanism of *Pg*DHAR and *Hs*CLIC1. Chemically modified *Pg*DHAR shows no activity while retaining the overall structure as confirmed by CD spectroscopy (Figure 3a, c). Chemical modification of *Hs*CLIC1:C24 with IAA and dimedone also resulted in complete loss of activity (Figure 3b, d), in concurrence with the loss of function mutation of Cys24 reported earlier [4]. Notably, all reported modifications carried out in plant DHARs are in concurrence with the loss-of-function mutation of N-terminal cysteine in the predicted active site [6, 14]. These results demonstrate the indispensable role of the first cysteine of the CXX[C/S] motif in DHARs and warrant the functional annotation of soluble *Hs*CLICs as DHA reductases.

**Figure 3.**
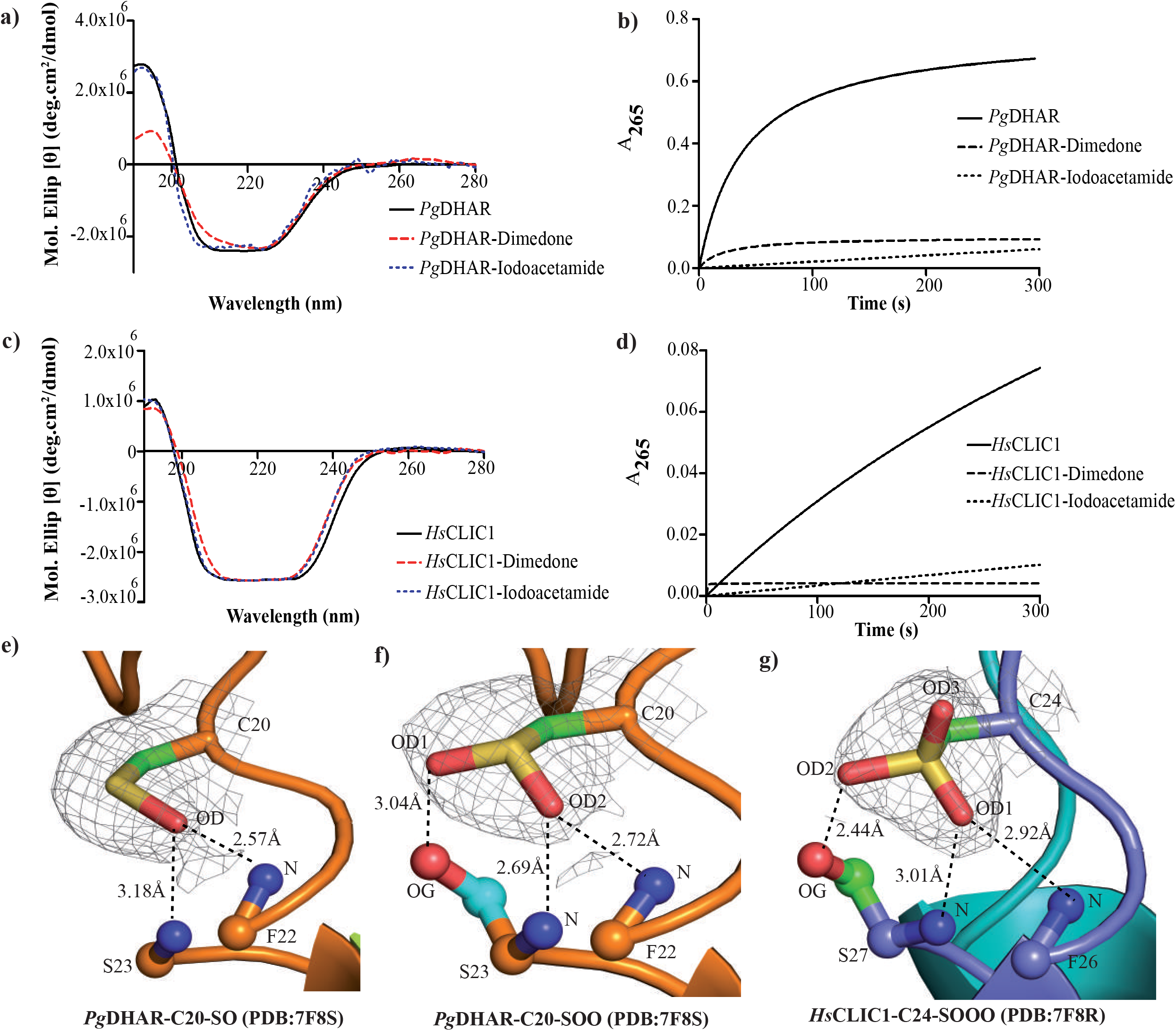
Chemical modification and crystallographic snapshots of reactive cysteine in *Pg*DHAR and *Hs*CLIC1 in various oxidation states. Effect of chemical modification on the overall structure of **(a)** *Pg*DHAR and **(c)** *Hs*CLIC1 as analysed using circular dichroism spectroscopy is shown. Corresponding effect on DHA reductase activity of **(b)** *Pg*DHAR and **(d)** *Hs*CLIC1 shown as time versus absorbance plot. Reactive cysteine captured in various redox states: **(e)** sulfenic, **(f)** sulfinic, and **(g)** sulfonic forms. Electron density contoured at 1σ (2Fo-Fc).

We then determined the crystal structures of *Pg*DHAR and *Hs*CLIC1 depicting the catalytic cysteine in various oxidation states: SO-sulfenic, SOO-sulfinic, SOOO-sulfonic forms (Figure 3e-g and Table 2). Of these, the sulfenic form is reverted to sulfhydryl by the GSH pool *in vivo*, unlike the sulfinic and sulfonic forms, which represent oxidatively damaged enzymes. These results demonstrate the indispensable role of the first cysteine of the CXX[C/S] motif in DHARs and warrant the functional annotation of soluble *Hs*CLICs as DHA reductases. Further, we investigated the role of additional residues present in the active site of these enzymes.

**Table 2.** X-ray data collection and refinement statistics.

|  | *Hs*CLIC1-C24-SOOO<br>(PDB: 7F8R) | *Pg*DHAR-C20-SO (A monomer)<br>*Pg*DHAR-C20-SOO (B monomer)<br>(PDB: 7F8S) |
| --- | --- | --- |
| **Data collection** |  |  |
| Space group | P 1 2<sub>1</sub> 1 | P 4<sub>3</sub> 2<sub>1</sub> 2 |
| **Cell dimensions** |  |  |
| *a*, *b*, *c* (Å) | 42.02, 70.66, 82.41 | 97.56, 97.56, 109.06 |
| $\alpha$, $\beta$, $\gamma$ (°) | 90, 90.13, 90 | 90, 90, 90 |
| Resolution (Å) | 42.06-2.51<br>(2.549-2.503) | 48.78-2.632<br>(2.726-2.632) |
| *R*<sub>sym</sub> or *R*<sub>merge</sub> | 0.062 (0.442) | 0.168 (0.728) |
| *I*/σ*I* | 16.2 (3.2) | 16.3 (3.9) |
| Completeness (%) | 97.2 (94.0) | 98.8 (97.2) |
| Redundancy | 4.1(4.0) | 14.7(13.6) |
| **Refinement** |  |  |
| Resolution (Å) | 2.51 | 2.63 |
| No. reflections | 16309 | 15997 |
| *R*<sub>work</sub>/*R*<sub>free</sub> | 0.185/0.235 | 0.207/0.250 |
| No. atoms | 3723 | 3372 |
| Protein | 3640 | 3248 |
| Ligand/ion | - | 45 |
| Water | 83 | 78 |
| B-factors<sup>‡</sup> (Å<sup>2</sup>) | 47.07 | 35.03 |
| Protein | 47.63 | 34.63 |
| Ligand/ion | - | 77.13 |
| Water | 40.52 | 27.33 |
| **R.m.s deviations** |  |  |
| Bond lengths (Å) | 0.016 | 0.019 |
| Bond angles (°) | 2.05 | 1.85 |

<sup>‡</sup> B-factor values shown are average of B-factors for respective atoms

### Role of H-site residues in DHA reductase activity of *Pg*DHAR and *Hs*CLICs

It is evident from electrostatic surface analysis (Figure 1d) that multiple conserved residues are present in the active site of these proteins. As members of the GST-omega family, both *Pg*DHAR and *Hs*CLICs have significant conservation across the G-site, where GSH binds. Hence, we investigated if any of the residue(s) surrounding the catalytic cysteine, but not directly associated with the G-site, are functionally important. Closer scrutiny of *Hs*CLIC1 with *Pg*DHAR shows that K8 and D19 in *Pg*DHAR and K13 of *Hs*CLIC1 are predicted to be in the ‘H-site’. Notably, these residues are highly conserved in DHARs and perhaps aid and confer substrate binding and specificity. Therefore, point mutagenesis was carried out to replace these residues with nonpolar amino acid alanine. Mutated *Hs*CLIC1 and *Pg*DHAR were expressed as soluble proteins in *E. coli* and purified to homogeneity. The kinetic constants of these mutants were determined and compared with respective native enzymes (Table 1). There was a dramatic reduction in the activity of mutants when compared to native *Pg*DHAR and *Hs*CLIC1. *Pg*DHAR-K8A shows a significant decrease in affinity for DHA (K_m_ DHA 1842μM (*Pg*DHAR-K8A) *vs* 159.2 μM (*Pg*DHAR)) without markedly affecting the K_m_ for GSH (K_m_ GSH 1885μM (*Pg*DHAR-K8A) *vs* 1477μM (*Pg*DHAR)). *Pg*DHAR-D19A did not have a pronounced effect on K_m_ for DHA (K_m_ DHA 245.7μM (*Pg*DHAR-D19A) *vs* 159.2μM (*Pg*DHAR)) however, affinity for GSH had significantly increased (K_m_ GSH 730.9 μM (*Pg*DHAR-D19A) *vs* 1477μM (*Pg*DHAR)). The corresponding K13A mutant of *Hs*CLIC1, similar to K8A mutant of *Pg*DHAR, shows a significant reduction in affinity to DHA (K_m_ DHA 4121 μM (*Hs*CLIC1-K13A) *vs* 2753μM (*Hs*CLIC1)). However, there was also a concomitant reduction in affinity to GSH (K_m_ GSH 7923 μM (*Hs*CLIC1-K13A) *vs* 2535μM (*Hs*CLIC1)) similar to *Pg*DHAR-K8A mutants. The specific activity and k_cat_ of *Hs*CLIC1 K13A mutant decreased 3.2 and 27-fold, respectively (specific activity 0.372 μM min^-1^mg^-1^; k_cat_ 0.012min^-1^), when compared to native enzyme (1.209 μM min^-1^mg^-1^; k_cat_ 0.325 min^-1^). Mutation of charged Lys8 and Asp19 residues in the H-site to Ala in *Pg*DHAR resulted in ∼96% and ∼99.1% reduction in specific activity, respectively, compared to native enzyme. Similarly K8A mutant of *Pto*DHAR2 shows 7.5% reduction in activity [14] and K11A mutant of *Cr*DHAR1 shows ∼54-32% reduction in activity when compared to native proteins [6]. Our data for *Pg*DHAR-K8A is in agreement with *Pto*DHAR-K8A and to a lesser extent *Cr*DHAR1-K11A (Supplementary Table 3), wherein the activity is not completely abolished. The complete loss of activity in *Pg*DHAR-D19A mutant correlates well with the previous data reported *Cr*DHAR1-D21N [6]. These reported mutations in H-site suggest a key role for these amino acids in DHA reductase activity.

### Implications of *Pg*DHAR in crop improvement

For this comparative study, we used a highly purified recombinant DHAR from *Pennisetum glaucum (L*.*) R. Br* (pearl millet). *P. glaucum* is a drought-resistant C4 plant from the Poaceae family, typically grown in semi-arid regions in India and Africa [45]. India is one of the largest producers of pearl millet in Asia and in the world [46]. Among C4 plants, pearl millet is comparatively more tolerant to drought than maize [47]. Multiple studies have reported the overall effect of overexpression of DHAR in plants whereas only a few have focused on the physiological relevance of AsA recycling by DHARs in drought-resistant varieties. Further, majority of the reported data on plant DHAR activity are inaccurate where quantification assays have been done using crude plant extracts that may have contaminating thioredoxins and glutaredoxins, which can also reduce DHA. Here, we have analysed the AsA recycling capacity of purified recombinant cytosolic DHAR from pearl millet. Comparative kinetic analysis with other plant DHARs reveals that *Pg*DHAR is the most efficient enzyme characterized to date. *Pg*DHAR shows ∼60 and 23-fold increased specific activity than that of *Zm*DHAR and *Os*DHAR, respectively. Increased DHAR activity is known to regulate the cellular AsA/DHA ratio which in turn will result in increased AsA pool [14, 15]. Out of synthesis, transportation, recycling and degradation, recycling is the primary mechanism by which plants maintain AsA homeostasis, as biosynthesis takes hours when compared to recycling [35]. In a recent transcriptome analysis, CAT1 (catalase1) and APX1 (ascorbate peroxidase1) transcript levels were found to be significantly higher in Pearl millet than Zea mays [47]. Higher AsA levels in turn would facilitate increased APX dependent reduction of H_2_O_2_ to water, therefore aiding pearl millet to withstand extreme climatic conditions such as drought. Thus, a detailed understanding of enzyme activity of various isoforms, paralogs and orthologs of plant DHARs may be useful for designing future transgenics aimed at improving drought resistance trait.

Further, it is predicted that there will be extreme drought conditions sub-setting in western hemisphere till 2099 [48] which will reduce the production of millets by at least 25% [49]. Improved drought-tolerant millet varieties will therefore be economically important in such geo-climatic conditions because they are expected to increase productivity and nutritional content. Hence, engineering *Pg*DHAR could improve the sustainability of normal millets and other food crops under very challenging climatic conditions.

### Implications for targeting Vitamin-C recycling *Hs*CLICs for anti-cancer therapy

Vitamin-C recycling is far more vital in humans as they can neither synthesize nor store AsA in higher quantities as plants do. Several distinct animals such as primates of the Haplorrhine order, some passeriform birds, bats and guinea pigs, also have lost the ability to synthesize AsA, and thus solely depend on dietary supply and recycling of AsA from DHA [50-52]. Whereas in plants the role of the DHAR as the primary enzyme involved in recycling AsA is well established, the enzyme(s) responsible for this activity in humans is not rigorously identified. Though predicted, the shared functional orthology of soluble plant DHARs and *Hs*CLICs was not clearly understood. Recent reports demonstrate, using a coupled enzyme assay, glutaredoxin-like oxidoreductase activity for purified *Hs*CLIC1 and 3, however, they lack kinetic and mechanistic insights [4, 21]. Our data clearly shows that both *Pg*DHAR and soluble *Hs*CLICs are mechanistically similar GSH-dependent DHA reductases, irrespective of their differences in kinetic characteristics. Both plant DHAR and *Hs*CLICs show a wide subcellular distribution with the former detected in cytosol, chloroplast, mitochondria and peroxisomes, and the latter in cytosol, nucleus and mitochondria [53, 54]. While *Pg*DHAR is upregulated under induced oxidative stress [32], *Hs*CLICs are significantly upregulated in multiple types of cancers. More so, cancer cells are under persistent oxidative stress [55, 56]. Thus, we speculate that overexpression of soluble enzymatic forms of *Hs*CLIC may be a survival strategy for cancer cells to escape elevated levels of ROS through increased ascorbate recycling. Further, the enzymatic activity of soluble vesicle-secreted *Hs*CLIC3 is implicated in invasion and metastasis [21]. As *Hs*CLICs have conserved G- and H-sites, involved in Vitamin-C recycling, both sites could be targeted for the development of targeted anti-cancer therapeutics.

Here, we have performed and compared the detailed kinetics of plant and human DHA reductases that recycle AsA; *Pg*DHAR and *Hs*CLIC1, 3 and 4, belonging to the GST-Omega family. Our data shows that DHAR from stress-adapted pearl millet is a suitable candidate for food crop improvement and the soluble enzymatic form of *Hs*CLICs 1, 3 and 4 could be targeted for novel anti-cancer therapy.

## Acknowledgements

Authors thank Murali Kaja, Emory Univ., for cDNA clones of *Hs*CLICs; Philip Board, The Australian National University, Canberra for the GST-O1 clone; VS Reddy for help with mass-spectrometry; CSIR for fellowship to BKD, DST-INSPIRE fellowship for SNS (IF150373) and DBT for post-doctoral fellowship to KP; Research in AA laboratory is funded by grants from Department of Biotechnology (BT/PR8766/BRB/10/1701/2018, BT/PR28080/BID/7/836/2018), Department of Science and Technology (EMR/2017005066) and ICGEB Core funds.

## Author contributions

BKD generated the expression constructs of native and mutant *Pg*DHAR and *Hs*CLICs, BKD and AK expressed, purified *Pg*DHAR and *Hs*CLICs and their mutants. BKD, KP, KG performed enzyme assays, biochemical and biophysical characterization. BKD, SNS, KG and KP analysed biochemical data. SK performed mass-spectrometry and analysed data. BKD, AK and SNS solved the crystal structures. MKR and MA provided *Pg*DHAR clones, materials and gave inputs for experiments and data analysis. BKD, AK, SNS and AA wrote the manuscript with inputs from all the authors.

## Data availability

The atomic coordinates and structure factors are available from Protein Data Bank under the accession codes 7F8R, 7F8S for *Hs*CLIC1-oxidised and *Pg*DHAR-oxidised structures, respectively.

## X-ray raw data

The raw X-ray diffraction data is available from Integrated Resource for Reproducibility in Macromolecular Crystallography (IRRMC) repository: https://proteindiffraction.org/

## Figure legends and Table legends

**Supplementary Table 1.** Structural and sequence identity of plant DHARs and *Hs*CLICs.

**Supplementary Table 2.** Enzyme kinetic data reported for various plant DHARs.

**Supplementary Table 3.** Kinetic data for other mammalian enzymes with DHA reductase activity.

**Supplementary Table 4.** MS/MS data for native and chemically modified *Pg*DHAR and *Hs*CLIC1.

**Supplementary Table 5.** Constructs, strains, and oligos used in this study.

---

**Supplementary Figure 1.**
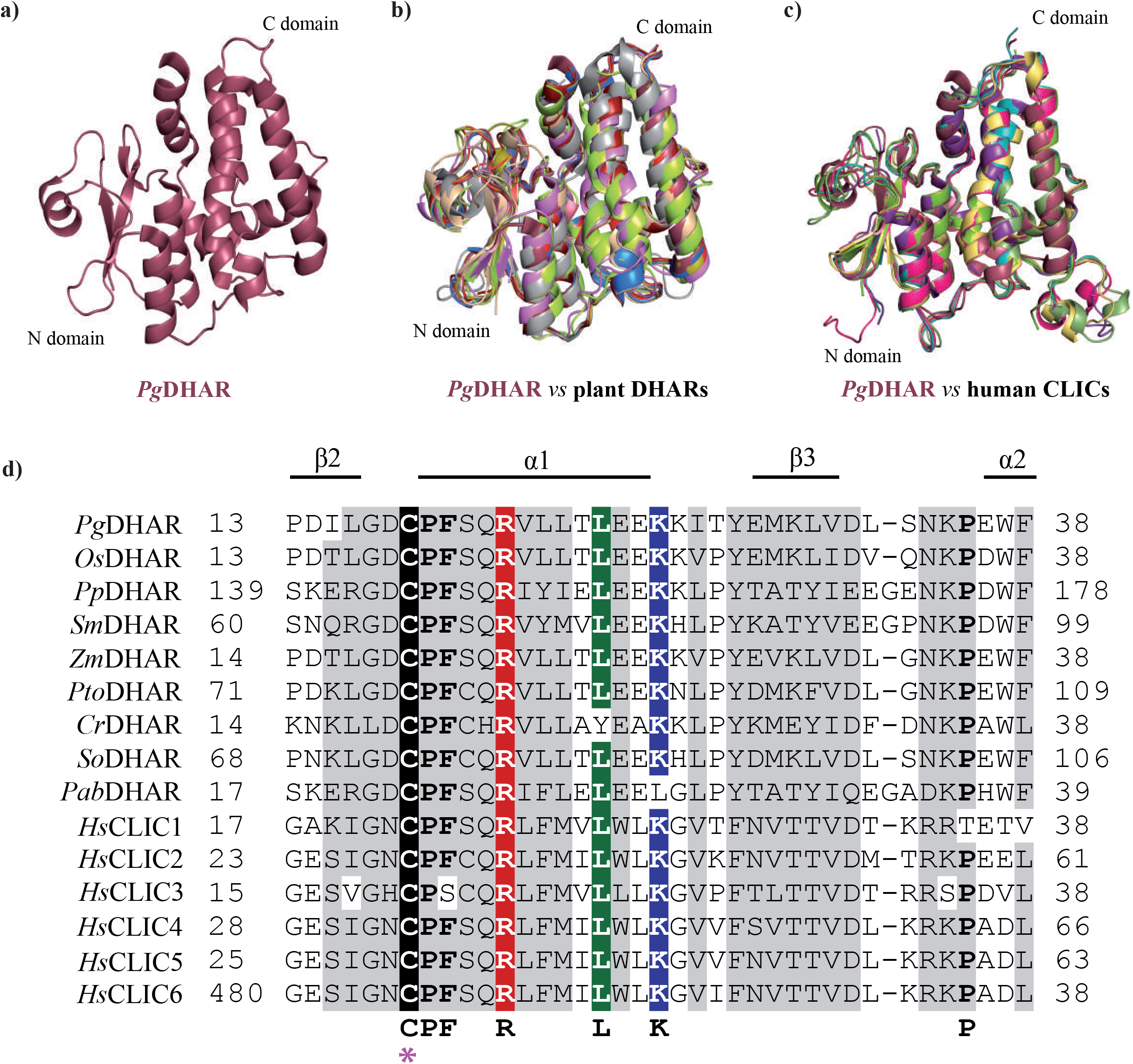
Structural and sequence comparison of plant and human DHA reductases. Structural superimposition of **(a)** *Pg*DHAR with **(b)** plant DHARs and **(c)** *Hs*CLICs1-6 with their corresponding structure guided sequence alignment are shown above. Proteins are colour coded as: *Pg*DHAR (5IQY) in raspberry, *Os*DHAR (5D9T) in marine, *Pp*DHAR (Swiss-Model) in tv_red, *Sm*DHAR (Swiss-Model) in olive, *Zm*DHAR (Swiss-Model) in firebrick, *Pto*DHAR (Swiss-Model) in wheat, *Cr*DHAR (5XFT) in gray70, *So*DHAR (Swiss-Model) in violet, *Pab*DHAR (Swiss-Model) in limon, *Hs*CLIC1 (1K0N) in split pea, *Hs*CLIC2 (2R4V) in violet purple, *Hs*CLIC3 (3FY7) in yellow orange, *Hs*CLIC4 (2D2Z) in teal, *Hs*CLIC5 (6Y2H) in hot pink, *Hs*CLIC6 (Swiss-Model) in green smudge. Species abbreviations are as follows: *Pg*: *Pennisetum glaucum, Os*: *Oryza sativa, Pp: Physcomitrella patens, Sm: Selaginella moellendorffii, Zm: Zea mays, Pt: Populus tomentosa, Cr: Chlamydomonas reinhardtii, So: Spinacia oleracea, Pab: Picea abies Hs*: *Homo sapiens*.

**Supplementary Figure 2.**
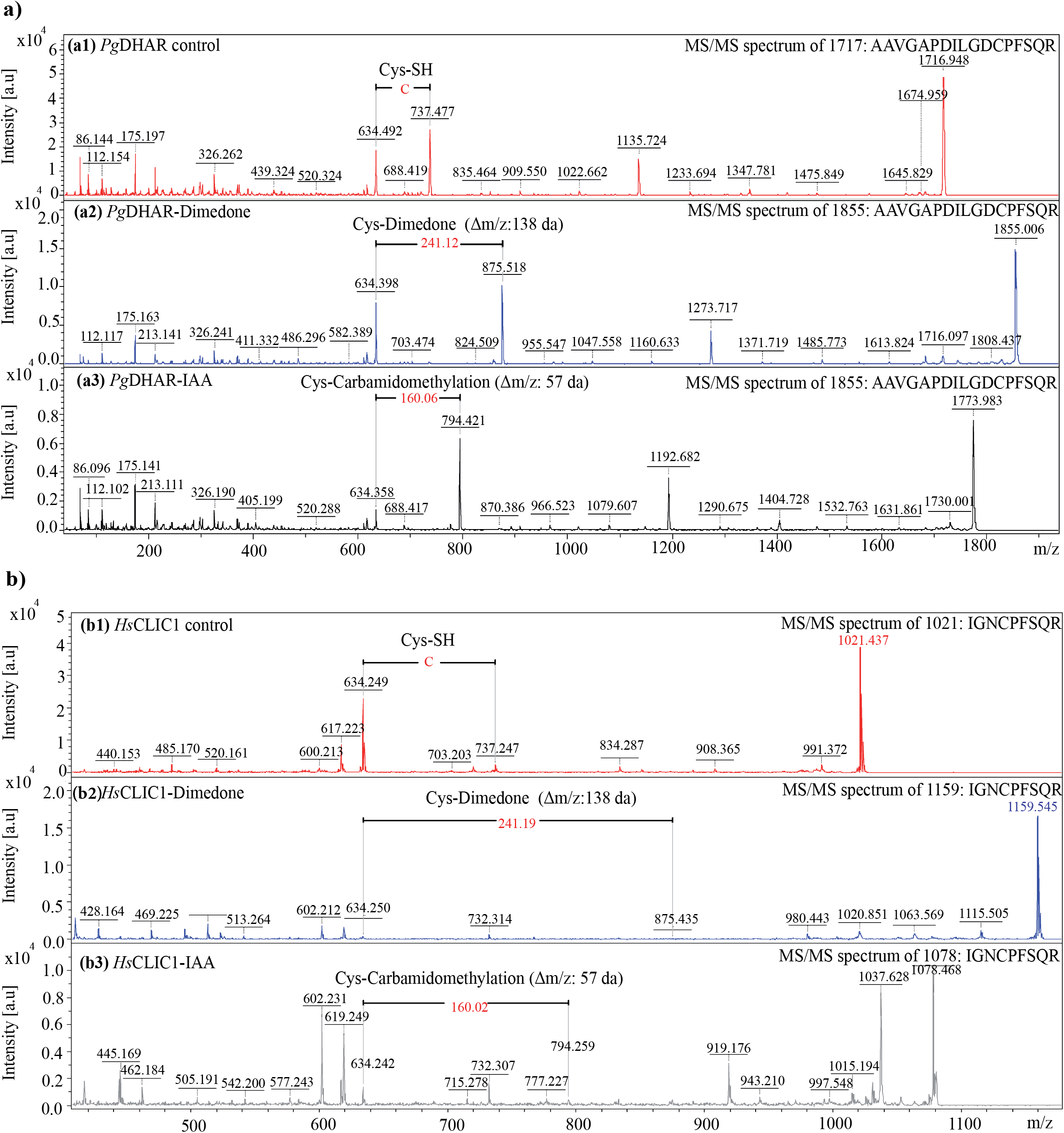
Mass-spectrometry analysis of catalytic cysteines. Tandem mass spectrum (MS/MS) of tryptic peptides-AAVGAPDILGDCPFSQR from **(a)** *Pg*DHAR and, IGNCPFSQR from **(b)** *Hs*CLIC1, showing catalytic cysteine (highlighted in yellow) in reduced - SH (a1,b1), dimedone modified - sulfenic -SO (a2,b2) and alkylated (a3,b3) forms.

**Supplementary Table 1.**
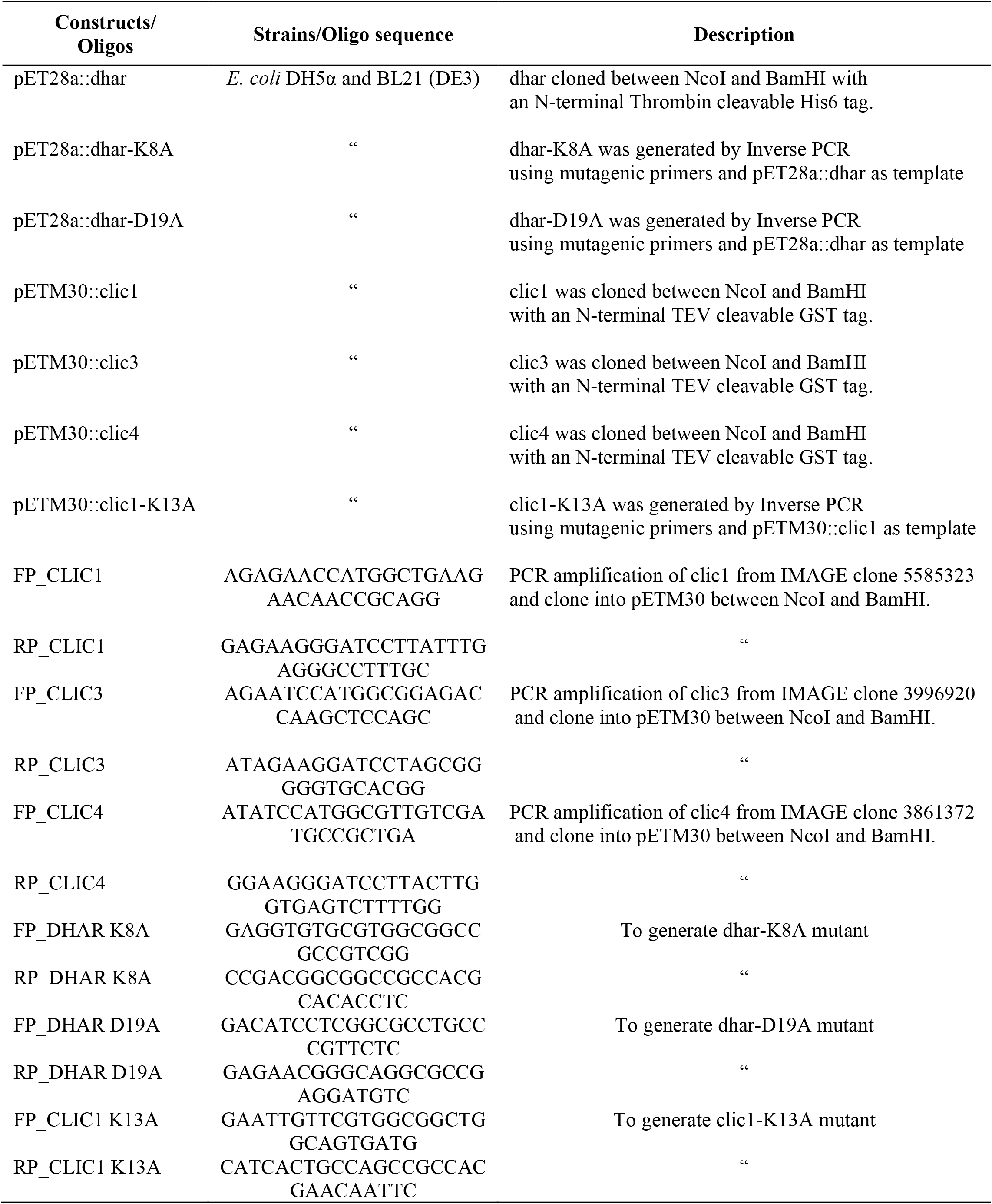
Constructs, strains, and oligos used in this study.

**Supplementary Table 2.** Structural and sequence identity of plant DHARs and *Hs*CLICs.

| **Protein** | **RMSD (Å)** | **%id** | **Z-Score** |
| --- | --- | --- | --- |
| *OsDHAR* | 0.5 | 84 | 35.6 |
| *PpDHAR* | 0.5 | 47 | 35.1 |
| *SmDHAR* | 0.6 | 47 | 35.0 |
| *ZmDHAR* | 0.7 | 90 | 35.4 |
| *PtoDHAR* | 1.5 | 62 | 30.9 |
| *CrDHAR* | 2.1 | 40 | 26.8 |
| *SoDHAR* | 2.2 | 59 | 23.2 |
| *PabDHAR* | 2.3 | 42 | 23.7 |
| *HsCLIC1* | 1.4 | 30 | 28.8 |
| *HsCLIC2* | 1.7 | 27 | 25.6 |
| *HsCLIC3* | 1.5 | 27 | 25.4 |
| *HsCLIC4* | 1.6 | 27 | 27.3 |
| *HsCLIC5* | 1.6 | 26 | 26.8 |
| *HsCLIC6*-(modelled protein) | 1.9 | 27 | 26.6 |

**Supplementary Table 3.**
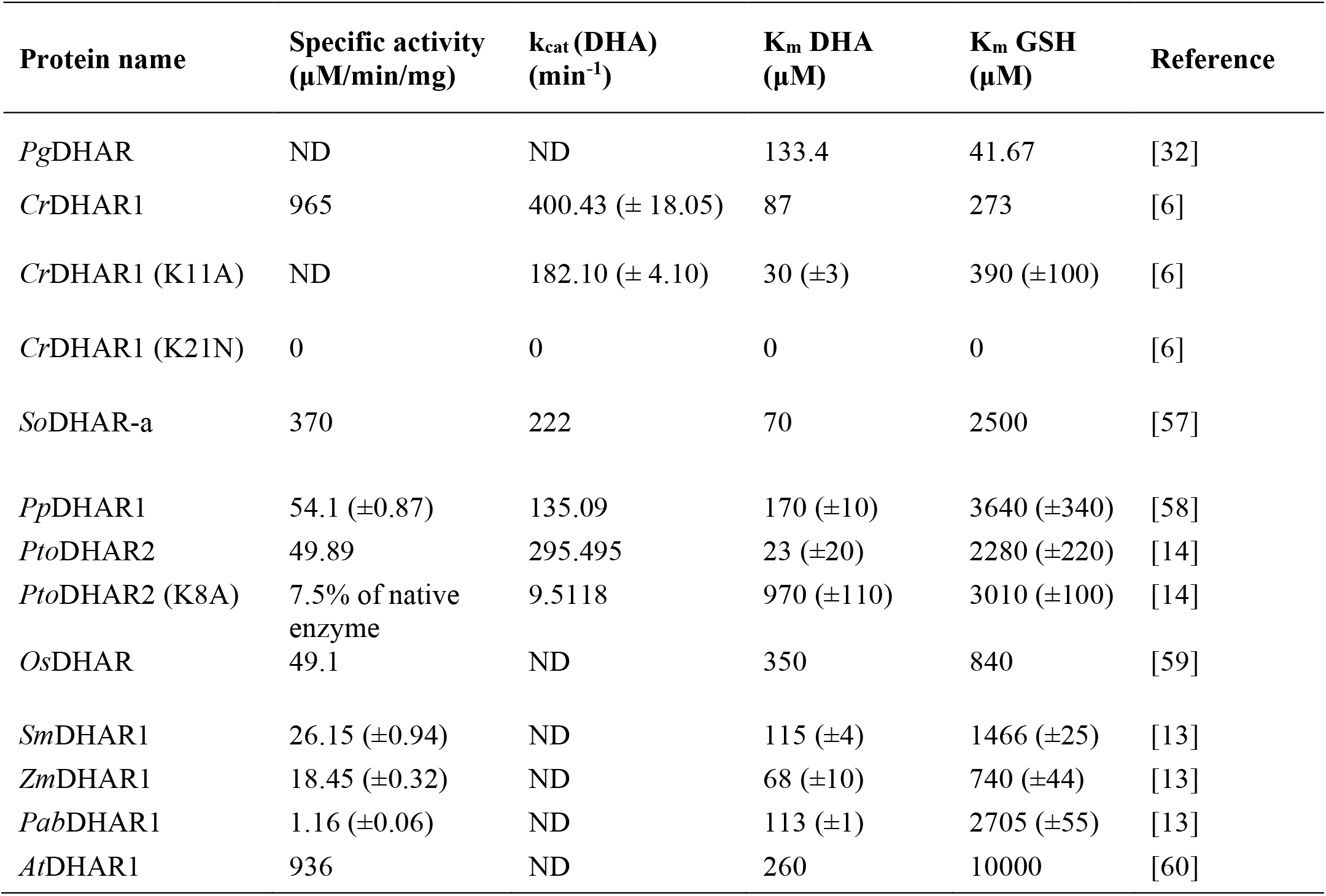
Enzyme kinetic data reported for various plant DHARs.

| Protein name | Specific activity<br>($\mu\text{M}/\text{min}/\text{mg}$) | $k_{\text{cat}}$ (DHA)<br>($\text{min}^{-1}$) | $K_{\text{m}}$ DHA<br>($\mu\text{M}$) | $K_{\text{m}}$ GSH<br>($\mu\text{M}$) | Reference |
| --- | --- | --- | --- | --- | --- |
| *Pg*DHAR | ND | ND | 133.4 | 41.67 | [32] |
| *Cr*DHAR1 | 965 | 400.43 (± 18.05) | 87 | 273 | [6] |
| *Cr*DHAR1 (K11A) | ND | 182.10 (± 4.10) | 30 (±3) | 390 (±100) | [6] |
| *Cr*DHAR1 (K21N) | 0 | 0 | 0 | 0 | [6] |
| *So*DHAR-a | 370 | 222 | 70 | 2500 | [57] |
| *Pp*DHAR1 | 54.1 (±0.87) | 135.09 | 170 (±10) | 3640 (±340) | [58] |
| *Pto*DHAR2 | 49.89 | 295.495 | 23 (±20) | 2280 (±220) | [14] |
| *Pto*DHAR2 (K8A) | 7.5% of native<br>enzyme | 9.5118 | 970 (±110) | 3010 (±100) | [14] |
| *Os*DHAR | 49.1 | ND | 350 | 840 | [59] |
| *Sm*DHAR1 | 26.15 (±0.94) | ND | 115 (±4) | 1466 (±25) | [13] |
| *Zm*DHAR1 | 18.45 (±0.32) | ND | 68 (±10) | 740 (±44) | [13] |
| *Pab*DHAR1 | 1.16 (±0.06) | ND | 113 (±1) | 2705 (±55) | [13] |
| *At*DHAR1 | 936 | ND | 260 | 10000 | [60] |

**Supplementary Table 4.** Kinetic data for other mammalian enzymes with DHA reductase activity.

| Protein name | Specific activity<br>($\mu\text{M}/\text{min}/\text{mg}$) | $k_{\text{cat}}$ (DHA)<br>($\text{min}^{-1}$) | $K_{\text{mDHA}}$<br>($\mu\text{M}$) | $K_{\text{m GSH}}$<br>($\mu\text{M}$) | Reference |
| --- | --- | --- | --- | --- | --- |
| *HsGSTO1-1* | 0.23<sup>a</sup> ($\pm 0.0160$)<br>0.14 ($\pm 0.013$) | ND | 138.2<sup>a</sup><br>($\pm 27.07$) | 2965<sup>a</sup> ($\pm 400.8$) | This study<br>[61] |
| *HsGSTO2-2* | 14.65 ($\pm 0.93$)<br>13.8 ($\pm 0.29$) | 0.66 ($\pm 0.03$) | 501 ($\pm 50$)<br>500 ($\pm 100$) | 4820 ($\pm 830$)<br>11400 ($\pm 5700$) | [39, 61] |
| Bovine PDI | 0.008 | ND | 1000 | 3900 | [62] |
| Erythrocyte<br>DHAR | 1.8 | 315 ($\pm 1$) | 210 ($\pm 60$) | 3500 ($\pm 300$) | [63] |
| *SmGSTO* | 0.2 | ND | 2300 | <4000 | [64] |
| Rat liver DHAR | 0.7645 | ND | 245 ($\pm 62$) | 2800 ($\pm 600$) | [65] |
| Pig GSTO1-1 | 0.1744 | ND | ND | ND | [66] |

Kinetic constants are the mean ± standard deviation of three or more separate experiments.

**Supplementary Table 5.** MS/MS data for native and chemically modified *Pg*DHAR and *Hs*CLIC1.

| Sample ID | DHAR- oxidized | DHAR- native | CLIC1- oxidized | CLIC1- native |
| --- | --- | --- | --- | --- |
| **Accession No.** | gi\|556824943 | gi\|556824943 | gi\|14251209 | gi\|14251209 |
| **Protein ID-**<br>**[Source]** | Dehydroascrobate<br>reductase<br>[*Cenchrus*<br>*americanus*] | Dehydroascrobate<br>reductase<br>[*Cenchrus*<br>*americanus*] | chloride<br>intracellular<br>channel protein 1<br>[*Homo sapiens*] | chloride<br>intracellular<br>channel protein 1<br>[*Homo sapiens*] |
| Theoretical Mol<br>wt. (da) | 23514 | 23514 | 27248 | 27248 |
| % Seq. Coverage | 26 | 7 | 27 | 2 |
| Theoretical pI | 5.98 | 5.98 | 5.09 | 5.09 |
| Observed m/z | 1481.9744 | 1716.9513 | 957.5638 | 1021.4315 |
|  | 1589.0186 |  | 1037.6415 |  |
|  | 1608.0198 |  | 1078.4769 |  |
|  | 1774.0433 |  | 1572.9301 |  |
|  |  |  | 2992.5467 |  |
| MS/MS- peptide<br>sequenced | K.LFHLQVALE<br>HFK.G | K.AAVGAPDIL<br>GDCPFSQR.V+<br>Cys (SH)* | R.YLSNAYAR.E | K.IGNCPFSQR.L<br>+ Cys (SH)* |
|  | K.LVDLSNKPE<br>WFLK.I |  | R.GFTIPEAFR.G |  |
|  | K.ALLDELQAL<br>DEHLK.A |  | K.IGNCPFSQR.L<br>+Carbamidometh<br>yl (C)* |  |
|  | K.AAVGAPDIL<br>GDCPFSQR.V+C<br>arbamidomethyl<br>(C)* |  | K.IEEFLEAVLC<br>PPR.Y |  |
|  |  |  | K.VLDNYLTSPL<br>PEEVDETSAD<br>EGVSQR.K |  |
| Individual Ion<br>score | 94 | 156 | 43 | 66 |
|  | 81 |  | 44 |  |
|  | 76 |  | 46 |  |
|  | 145 |  | 105 |  |
|  |  |  | 48 |  |
| MOWSE | 224 | 156 | 286 | 66 |

## References

1. Foyer CH & Halliwell B (1976) The presence of glutathione and glutathione reductase in chloroplasts: a proposed role in ascorbic acid metabolism. Planta 133, 21–25.

2. Foyer CH & Halliwell B (1977) Purification and properties of dehydroascorbate reductase from spinach leaves. Phytochemistry 16, 1347–1350.

3. Asada K (1999) The water-water cycle in chloroplasts: scavenging of active oxygens and dissipation of excess photons. Annual review of plant biology 50, 601–639.

4. Al Khamici H, Brown LJ, Hossain KR, Hudson AL, Sinclair-Burton AA, Ng JP, Daniel EL, Hare JE, Cornell BA, Curmi PM, et al. (2015) Members of the chloride intracellular ion channel protein family demonstrate glutaredoxin-like enzymatic activity. PLoS One 10, e115699, doi: 10.1371/journal.pone.0115699.

5. Lallement PA, Roret T, Tsan P, Gualberto JM, Girardet JM, Didierjean C, Rouhier N & Hecker A (2016) Insights into ascorbate regeneration in plants: investigating the redox and structural properties of dehydroascorbate reductases from Populus trichocarpa. Biochem J 473, 717–731, doi: 10.1042/BJ20151147.

6. Chang HY, Lin ST, Ko TP, Wu SM, Lin TH, Chang YC, Huang KF & Lee TM (2017) Enzymatic characterization and crystal structure analysis of Chlamydomonas reinhardtii dehydroascorbate reductase and their implications for oxidative stress. Plant Physiol Biochem 120, 144–155, doi: 10.1016/j.plaphy.2017.09.026.

7. Yin L, Wang S, Eltayeb AE, Uddin MI, Yamamoto Y, Tsuji W, Takeuchi Y & Tanaka K (2010) Overexpression of dehydroascorbate reductase, but not monodehydroascorbate reductase, confers tolerance to aluminum stress in transgenic tobacco. Planta 231, 609–621, doi: 10.1007/s00425-009-1075-3.

8. Noshi M, Yamada H, Hatanaka R, Tanabe N, Tamoi M & Shigeoka S (2017) Arabidopsis dehydroascorbate reductase 1 and 2 modulate redox states of ascorbate-glutathione cycle in the cytosol in response to photooxidative stress. Biosci Biotechnol Biochem 81, 523–533, doi: 10.1080/09168451.2016.1256759.

9. Smirnoff N & Wheeler GL (2000) Ascorbic acid in plants: biosynthesis and function. Crit Rev Biochem Mol Biol 35, 291–314, doi: 10.1080/10409230008984166.

10. Shimaoka T, Miyake C & Yokota A (2003) Mechanism of the reaction catalyzed by dehydroascorbate reductase from spinach chloroplasts. Eur J Biochem 270, 921–928, doi: 10.1046/j.1432-1033.2003.03452.x.

11. Do H, Kim IS, Jeon BW, Lee CW, Park AK, Wi AR, Shin SC, Park H, Kim YS, Yoon HS, et al. (2016) Structural understanding of the recycling of oxidized ascorbate by dehydroascorbate reductase (OsDHAR) from Oryza sativa L. japonica. Sci Rep 6, 19498, doi: 10.1038/srep19498.

12. Bartoli CG, Guiamet JJ, Kiddle G, Pastori GM, Di Cagno R, Theodoulou FL & Foyer CH (2005) Ascorbate content of wheat leaves is not determined by maximal l‐galactono‐1, 4‐lactone dehydrogenase (GalLDH) activity under drought stress. Plant, Cell & Environment 28, 1073–1081.

13. Zhang YJ, Wang W, Yang HL, Li Y, Kang XY, Wang XR & Yang ZL (2015) Molecular Properties and Functional Divergence of the Dehydroascorbate Reductase Gene Family in Lower and Higher Plants. PLoS One 10, e0145038, doi: 10.1371/journal.pone.0145038.

14. Tang ZX & Yang HL (2013) Functional divergence and catalytic properties of dehydroascorbate reductase family proteins from Populus tomentosa. Mol Biol Rep 40, 5105–5114, doi: 10.1007/s11033-013-2612-5.

15. Kim Y-S, Kim I-S, Bae M-J, Choe Y-H, Kim Y-H, Park H-M, Kang H-G & Yoon H-S (2013) Homologous expression of cytosolic dehydroascorbate reductase increases grain yield and biomass under paddy field conditions in transgenic rice (Oryza sativa L. japonica). Planta 237, 1613–1625.

16. Littler DR, Harrop SJ, Goodchild SC, Phang JM, Mynott AV, Jiang L, Valenzuela SM, Mazzanti M, Brown LJ, Breit SN, et al. (2010) The enigma of the CLIC proteins: Ion channels, redox proteins, enzymes, scaffolding proteins? FEBS Lett 584, 2093–2101, doi: 10.1016/j.febslet.2010.01.027.

17. Berry KL, Bulow HE, Hall DH & Hobert O (2003) A C. elegans CLIC-like protein required for intracellular tube formation and maintenance. Science 302, 2134–2137, doi: 10.1126/science.1087667.

18. Suh KS, Mutoh M, Gerdes M, Crutchley JM, Mutoh T, Edwards LE, Dumont RA, Sodha P, Cheng C, Glick A, et al. (2005) Antisense suppression of the chloride intracellular channel family induces apoptosis, enhances tumor necrosis factor {alpha}-induced apoptosis, and inhibits tumor growth. Cancer Res 65, 562–571.

19. Valenzuela SM, Mazzanti M, Tonini R, Qiu MR, Warton K, Musgrove EA, Campbell TJ & Breit SN (2000) The nuclear chloride ion channel NCC27 is involved in regulation of the cell cycle. J Physiol 529 Pt 3, 541–552.

20. Averaimo S, Milton RH, Duchen MR & Mazzanti M (2010) Chloride intracellular channel 1 (CLIC1): Sensor and effector during oxidative stress. FEBS Lett 584, 2076–2084, doi: 10.1016/j.febslet.2010.02.073.

21. Hernandez-Fernaud JR, Ruengeler E, Casazza A, Neilson LJ, Pulleine E, Santi A, Ismail S, Lilla S, Dhayade S & MacPherson IR (2017) Secreted CLIC3 drives cancer progression through its glutathione-dependent oxidoreductase activity. Nature communications 8, 1–17.

22. Kumar S, Kumar K, Pandey P, Rajamani V, Padmalatha KV, Dhandapani G, Kanakachari M, Leelavathi S, Kumar PA & Reddy VS (2013) Glycoproteome of elongating cotton fiber cells. Mol Cell Proteomics 12, 3677–3689, doi: 10.1074/mcp.M113.030726.

23. Otwinowski Z & Minor W (1997) [20] Processing of X-ray diffraction data collected in oscillation mode. Methods in enzymology 276, 307–326.

24. Vonrhein C, Flensburg C, Keller P, Sharff A, Smart O, Paciorek W, Womack T & Bricogne G (2011) Data processing and analysis with the autoPROC toolbox. Acta Crystallogr D Biol Crystallogr 67, 293–302, doi: 10.1107/S0907444911007773.

25. Long F, Vagin AA, Young P & Murshudov GN (2008) BALBES: a molecular-replacement pipeline. Acta Crystallographica Section D: Biological Crystallography 64, 125–132.

26. Adams PD, Afonine PV, Bunkóczi G, Chen VB, Davis IW, Echols N, Headd JJ, Hung L-W, Kapral GJ & Grosse-Kunstleve RW (2010) PHENIX: a comprehensive Python-based system for macromolecular structure solution. Acta Crystallographica Section D: Biological Crystallography 66, 213–221.

27. Emsley P, Lohkamp B, Scott WG & Cowtan K (2010) Features and development of Coot. Acta Crystallographica Section D: Biological Crystallography 66, 486–501.

28. Vagin AA, Steiner RA, Lebedev AA, Potterton L, McNicholas S, Long F & Murshudov GN (2004) REFMAC5 dictionary: organization of prior chemical knowledge and guidelines for its use. Acta Crystallographica Section D: Biological Crystallography 60, 2184–2195.

29. Krishna Das B, Kumar A, Maindola P, Mahanty S, Jain SK, Reddy MK & Arockiasamy A (2016) Non-native ligands define the active site of Pennisetum glaucum (L.) R. Br dehydroascorbate reductase. Biochem Biophys Res Commun 473, 1152–1157, doi: 10.1016/j.bbrc.2016.04.031.

30. Oakley A (2011) Glutathione transferases: a structural perspective. Drug Metab Rev 43, 138–151, doi: 10.3109/03602532.2011.558093.

31. Frova C (2006) Glutathione transferases in the genomics era: new insights and perspectives. Biomol Eng 23, 149–169, doi: 10.1016/j.bioeng.2006.05.020.

32. Pandey P, Achary VM, Kalasamudramu V, Mahanty S, Reddy GM & Reddy MK (2014) Molecular and biochemical characterization of dehydroascorbate reductase from a stress adapted C4 plant, pearl millet [Pennisetum glaucum (L.) R. Br]. Plant Cell Rep 33, 435–445, doi: 10.1007/s00299-013-1544-9.

33. Dolinsky TJ, Nielsen JE, McCammon JA & Baker NA (2004) PDB2PQR: an automated pipeline for the setup of Poisson-Boltzmann electrostatics calculations. Nucleic Acids Res 32, W665–667, doi: 10.1093/nar/gkh381.

34. Chun JA, Lee WH, Han MO, Lee JW, Yi YB, Goo YM, Lee SW, Bae SC, Cho KJ & Chung CH (2007) Molecular and biochemical characterizations of dehydroascorbate reductase from sesame (Sesamum indicum L.) hairy root cultures. J Agric Food Chem 55, 6067–6073, doi: 10.1021/jf070946t.

35. Bartoli CG, Gomez F, Gergoff G, Guiamet JJ & Puntarulo S (2005) Up-regulation of the mitochondrial alternative oxidase pathway enhances photosynthetic electron transport under drought conditions. J Exp Bot 56, 1269–1276, doi: 10.1093/jxb/eri111.

36. Shimaoka T, Yokota A & Miyake C (2000) Purification and characterization of chloroplast dehydroascorbate reductase from spinach leaves. Plant Cell Physiol 41, 1110–1118, doi: 10.1093/pcp/pcd035.

37. Jung C-H & Wells WW (1998) Spontaneous Conversion ofl-Dehydroascorbic Acid tol-Ascorbic Acid andl-Erythroascorbic Acid. Archives of biochemistry and biophysics 355, 9–14, doi: 10.1006/abbi.1998.0713.

38. Schomburg I, Chang A & Schomburg D (2002) BRENDA, enzyme data and metabolic information. Nucleic Acids Research 30, 47–49, doi: 10.1093/nar/30.1.47.

39. Zhou H, Brock J, Liu D, Board PG & Oakley AJ (2012) Structural insights into the dehydroascorbate reductase activity of human omega-class glutathione transferases. J Mol Biol 420, 190–203, doi: 10.1016/j.jmb.2012.04.014.

40. Foyer CH & Noctor G (2005) Redox homeostasis and antioxidant signaling: a metabolic interface between stress perception and physiological responses. The Plant Cell 17, 1866–1875.

41. Sies H (2014) Role of metabolic H2O2 generation: redox signaling and oxidative stress. Journal of Biological Chemistry 289, 8735–8741.

42. Washburn MP & Wells WW (1999) The catalytic mechanism of the glutathione-dependent dehydroascorbate reductase activity of thioltransferase (glutaredoxin). Biochemistry 38, 268–274.

43. Marchand CH, Fermani S, Rossi J, Gurrieri L, Tedesco D, Henri J, Sparla F, Trost P, Lemaire SD & Zaffagnini M (2019) Structural and Biochemical Insights into the Reactivity of Thioredoxin h1 from Chlamydomonas reinhardtii. Antioxidants 8, 10.

44. Arts IS, Vertommen D, Baldin F, Laloux G & Collet J-F (2016) Comprehensively characterizing the thioredoxin interactome in vivo highlights the central role played by this ubiquitous oxidoreductase in redox control. Molecular & Cellular Proteomics 15, 2125–2140.

45. ICAR-AICRP on Pearl millet (2018) AICRP on Pearl Millet, Jodhpur, India. In.

46. IndexMundi (2020) Millet Production rate by Country in 1000 MT. In.

47. Iwuala E, Odjegba V, Ajiboye A, Umebese C, Sharma V & Alam A (2019) Effect of drought stress on the expression of genes linked to antioxidant enzymatic activity in landraces of Zea mays L. and Pennisetum glaucum (L.) R. Br. Plant Physiology Reports 24, 422–433, doi: 10.1007/s40502-019-00460-0.

48. Dai A (2013) Increasing drought under global warming in observations and models. Nature Climate Change 3, 52–58, doi: 10.1038/nclimate1633.

49. Choudhary M, Jayanand & Padaria JC (2015) Transcriptional profiling in pearl millet (Pennisetum glaucum L.R. Br.) for identification of differentially expressed drought responsive genes. Physiol Mol Biol Plants 21, 187–196, doi: 10.1007/s12298-015-0287-1.

50. Birney EC, Jenness R & Ayaz KM (1976) Inability of bats to synthesise L-ascorbic acid. Nature 260, 626–628, doi: 10.1038/260626a0.

51. Chaudhuri CR & Chatterjee IB (1969) L-ascorbic acid synthesis in birds: phylogenetic trend. Science 164, 435–436, doi: 10.1126/science.164.3878.435.

52. Montel-Hagen A, Kinet S, Manel N, Mongellaz C, Prohaska R, Battini JL, Delaunay J, Sitbon M & Taylor N (2008) Erythrocyte Glut1 triggers dehydroascorbic acid uptake in mammals unable to synthesize vitamin C. Cell 132, 1039–1048, doi: 10.1016/j.cell.2008.01.042.

53. Chen Z & Gallie DR (2006) Dehydroascorbate reductase affects leaf growth, development, and function. Plant Physiol 142, 775–787, doi: 10.1104/pp.106.085506.

54. Ulmasov B, Bruno J, Woost PG & Edwards JC (2007) Tissue and subcellular distribution of CLIC1. BMC Cell Biol 8, 8, doi: 10.1186/1471-2121-8-8.

55. Cairns RA, Harris IS & Mak TW (2011) Regulation of cancer cell metabolism. Nat Rev Cancer 11, 85–95, doi: 10.1038/nrc2981.

56. Toyokuni S, Okamoto K, Yodoi J & Hiai H (1995) Persistent oxidative stress in cancer. FEBS Lett 358, 1–3, doi: 10.1016/0014-5793(94)01368-b.

57. Hossain MA & Asada K (1984) Purification of Dehydroascorbate Reductase from Spinach and Its Characterization as a Thiol Enzyme. Plant and Cell Physiology 25, 85–92, doi: 10.1093/oxfordjournals.pcp.a076700.

58. Liu YJ, Han XM, Ren LL, Yang HL & Zeng QY (2013) Functional divergence of the glutathione S-transferase supergene family in Physcomitrella patens reveals complex patterns of large gene family evolution in land plants. Plant Physiol 161, 773–786, doi: 10.1104/pp.112.205815.

59. Kato Y, Urano Ji, Maki Y & Ushimaru T (1997) Purification and Characterization of Dehydroascorbate Reductase from Rice. Plant and Cell Physiology 38, 173–178, doi: 10.1093/oxfordjournals.pcp.a029149.

60. Dixon DP, Hawkins T, Hussey PJ & Edwards R (2009) Enzyme activities and subcellular localization of members of the Arabidopsis glutathione transferase superfamily. J Exp Bot 60, 1207–1218, doi: 10.1093/jxb/ern365.

61. Schmuck EM, Board PG, Whitbread AK, Tetlow N, Cavanaugh JA, Blackburn AC & Masoumi A (2005) Characterization of the monomethylarsonate reductase and dehydroascorbate reductase activities of Omega class glutathione transferase variants: implications for arsenic metabolism and the age-at-onset of Alzheimer’s and Parkinson’s diseases. Pharmacogenetics and Genomics 15, 493–501.

62. Wells WW, Xu DP, Yang Y & Rocque PA (1990) Mammalian thioltransferase (glutaredoxin) and protein disulfide isomerase have dehydroascorbate reductase activity. Journal of Biological Chemistry 265, 15361–15364.

63. Xu DP, Washburn MP, Sun GP & Wells WW (1996) Purification and characterization of a glutathione dependent dehydroascorbate reductase from human erythrocytes. Biochemical and biophysical research communications 221, 117–121.

64. Girardini J, Amirante A, Zemzoumi K & Serra E (2002) Characterization of an omega‐class glutathione S‐transferase from Schistosoma mansoni with glutaredoxin‐like dehydroascorbate reductase and thiol transferase activities. European journal of biochemistry 269, 5512–5521.

65. Maellaro E, Del Bello B, Sugherini L, Santucci A, Comporti M & Casini A (1994) Purification and characterization of glutathione-dependent dehydroascorbate reductase from rat liver. Biochemical Journal 301, 471–476.

66. Rouimi P, Anglade P, Benzekri A, Costet P, Debrauwer L, Pineau T & Tulliez J (2001) Purification and characterization of a glutathione S-transferase Omega in pig: evidence for two distinct organ-specific transcripts. Biochemical Journal 358, 257–262.

